# Keratinocyte-derived paracrine factors regulate stress response of melanocytes to UVB

**DOI:** 10.1101/2023.01.13.523939

**Authors:** Saowanee Jeayeng, Malinee Saelim, Phetthinee Muanjumpon, Pongsakorn Buraphat, Potjanee Kanchanapiboon, Somponnat Sampattavanich, Uraiwan Panich

## Abstract

The skin microenvironment created by keratinocytes (KC) influences stress responses of melanocytes (MC) to UVB insult. Here, we investigated paracrine factors involved in the regulatory role of microenvironment created by KC in UVB-mediated MC responses using RNA sequencing analysis as well as *in vitro* and *in vivo* models. RNA-Seq showed that G-CSF and CCL20 genes were highly upregulated in UVB-irradiated KC and their levels best correlated with paracrine protective effects of KC on stress responses of MC to UVB. Recombinant G-CSF and CCL20 treatment revealed the strongest modulatory effects on UVB-induced MC responses by mitigating apoptosis and ROS formation and upregulating tyrosinase and tyrosinase-related protein-1 (TRP-1) involved in the melanogenic pathway. A similar correlation between G-CSF and CCL20 expression in KC and the tyrosinase level in MC was also observed in the UVB-irradiated mouse skin. Our study reports for the first time that G-CSF and CCL20 might play a regulatory role in the KC’s paracrine effects on UVB-mediated MC damage and also provides translational insights for the development of biomarkers for predicting susceptibility to photodamage.

**Graphical abstract:** 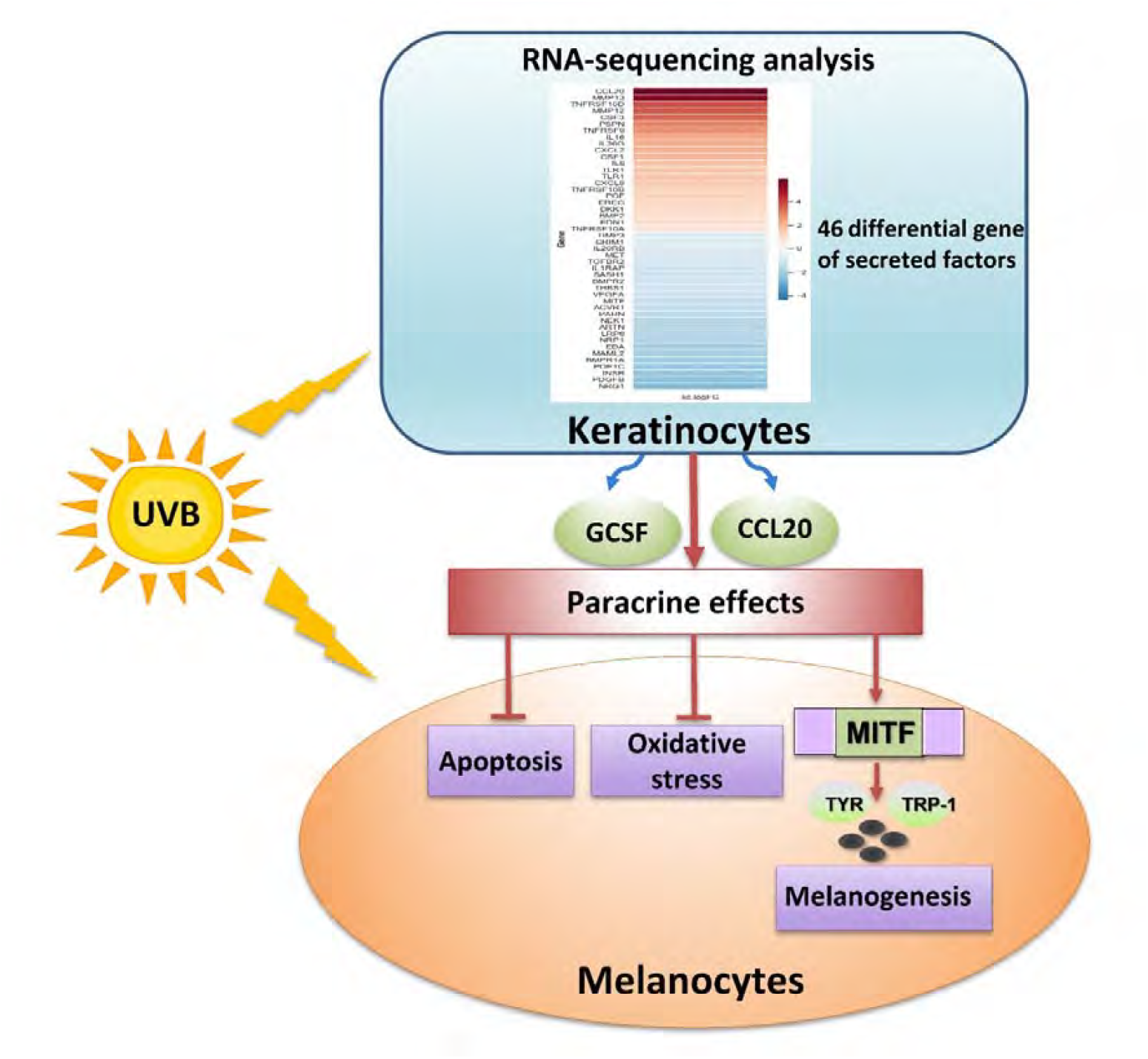

## Introduction

Accumulative exposure to ultraviolet radiation (UVR) including UVB is the primary factor causing cutaneous photodamage and photocarcinogenesis (Agar & Young, 2005; Brozyna *et al*, 2007; Jeayeng *et al*, 2017). Epidermal cells including keratinocytes (KC) and melanocytes (MC) act as sensors of environmental stresses affecting the skin. KC, representing at least 90% of the epidermal cells, are the first line of defense against external insults including UVR (Choi *et al*, 2010). The microenvironment created by KC differentially influences homeostasis, function (including melanogenesis) and phenotype of the adjacent MC in response to UVR. Chronic exposure of the skin to UVB can mediate disrupted homeostasis of MC and cause stress responses including apoptosis (Brozyna *et al*, 2007), DNA damage (Premi *et al*, 2015) and oxidative stress (Chakraborty *et al*, 1999), ultimately contributing towards the development of malignant melanoma (Sample A and He YY, 2018). MC homeostasis is influenced by paracrine communication via several soluble factors such as growth factors, cytokines and hormones secreted from the neighboring KC (Choi *et al*, 2010). From prior work, KC-derived paracrine factors that can influence the MC response to UVB include endothelin-1 (ET-1), proopiomelanocortin (POMC) derived peptides [such as α-melanocyte-stimulating hormone (α-MSH) and adrenocorticotropic hormone (ACTH)] and corticotrophin-releasing hormone (CRH) (Bohm *et al*, 2005; Hyter *et al*, 2013; Kadekaro *et al*, 2005; Slominski *et al*, 2000; Slominski *et al*, 2013; Slominski *et al*, 2005; Jeayeng *et al*, 2017). Skin exposure to UVR also promotes KC to secrete several pro-inflammatory cytokines, e.g., IL-1β, IL-6 and tumor necrosis factor (TNF)-alpha, that can also exert physiological and biological effects on MC (Moretti *et al*, 2009). Finally, melanogenesis (pigmentation) is a vital adaptive protective response against further DNA damage induced by UVR (Nguyen *et al*, 2019). Upon UVB exposure, KC-derived paracrine factors (e.g., α-MSH) can control melanogenesis and affect the survival and the differentiation of MC vi Mitf, which is regulated by MAPK signaling (Juemin *et al*, 2022). Our previous work also showed that KC exerted a paracrine protective effect on UVB-mediated DNA damage, apoptosis and inflammatory responses in association with MAPK pathways in MC (Jeayeng *et al*, 2017).

Since crosstalk between KC’s microenvironment and MC plays a critical role in regulating skin homeostasis, identification of the key KC-derived paracrine factors will be useful for the development of biomarkers for predicting UVR sensitivity and the risk of photodamage and can lead to the development of a novel and effective strategy for skin cancer prevention.

In this study, we were interested in identifying KC-derived paracrine factors that can protect MC from photodamage. By examining the secreted factors from different epidermal cell sources that showed correlated UVB-protective effects, we could identify the candidate paracrine factors from transcriptomic profiling. We then confirmed the identity of paracrine factors that exhibit the strongest protective effect against UVB in MC using recombinant proteins. Finally, the proposed paracrine interactions between KC and MC were validated *in vivo* using a mouse skin model.

## Results

### The paracrine protective effects on UVB-induced apoptosis, ROS formation and melanogenesis in MC

We were interested in identifying the KC-derived paracrine factors that underlie protective effects in UVB-irradiated MC. We chose five epidermal cell sources (KC, HaCaT, HDF, NIHT3T, and A431 cells) to assess their differential influences on UVB-mediated response in MC. We reasoned that the expression of key paracrine factors from KC must be proportional to the observed UVB-mediated phenotypic changes in MC. At first, the paracrine release was stimulated by exposing these five dermal cell sources to UVB irradiation (125 mJ/cm^2^) for 12 hours. Conditioned media (CM) from these five dermal cell sources were collected and used to incubate MC for 2 hours prior to UVB irradiation. Various UVB-induced phenotypic changes in MC including apoptosis, oxidative stress and melanogenesis were compared when MC cells were cultured with or without CM. Without CM, UVB irradiation markedly stimulated caspase3 activation (37.34 ± 6% of positive-stained cells) (Fig 1A) and oxidant formation (71.08 ± 1% of positive-stained cells) (Fig 1B) in MC alone. Moreover, UVB irradiation resulted in a significant increase in melanin content (125.11 ± 18% of control) (Fig 1C) and tyrosinase activity (120.66 ± 24.9% of control) (Fig 1D) in MC. With CM, we observed a substantial reduction of UVB-mediated active caspase-3 and ROS production in MC at varying degrees from different CM sources: KC (14.36 ± 2.1% and 41 ± 1%), HaCaT (18.4 ± 2.2% and 48.56 ± 2.3%), HDF (22.33 ± 5.2% and 53.25 ± 1.9%) and NIH 3T3 (25.16 ± 3% and 58.68 ± 1.6%) compared to untreated MC (37.34 ± 6% and 71.08 ± 1%)(Fig 1A and B). Furthermore, following UVB exposure, increases in melanin content and tyrosinase activity were observed in MC that were pre-incubated with different CM from different cell sources: KC (171.8 ± 11.4% and 177.74 ± 12.8%), HaCaT (254.52± 11.4% and 160.08 ± 13.4%), HDF (140.39 ± 11.9% and 157.91 ± 8.4%), NIH3T3 (135.86 ± 14.4% and 144.53 ± 23.6%) compared to untreated MC (125.11 ± 18% and 120.66 ± 24.9%) (Fig 1C and D). Our results demonstrated that CM from different UVB-irradiated cell types could indeed protect MC from the UVB-mediated stress responses in MC but at a different strength. When we comprehensively compared the UVB-protective effect across different dermal cells, we found that CM collected from skin cells including KC, HaCaT, HDF, NIH 3T3 except A431 cells were able to significantly suppress caspase-3 activation and ROS formation as well as increase tyrosinase activity and melanin level in UVB-irradiated MC (Fig 1E). Specifically, KC appeared to exert the strongest paracrine modulatory effects on UVB-mediated stress responses including photodamage and melanogenesis. To confirm these results, we attempted to assess the paracrine modulatory effects with altering cell density for CM collection. The different cell plating densities of 1:5 (MC:KC-CM) and 1:10 (MC:KC-CM) significantly provided paracrine protective effects against apoptosis of MC following UVB exposure, indicating that the protective effects from KC-derived CM against MC damage were dose-dependent. We chose the different cell plating densities (1:5; MC:KC-CM) for our further experiments since this condition created the strongest modulatory effect against stress response in UVB-irradiated MC (Appendix Fig S1).

**Figure 1.**
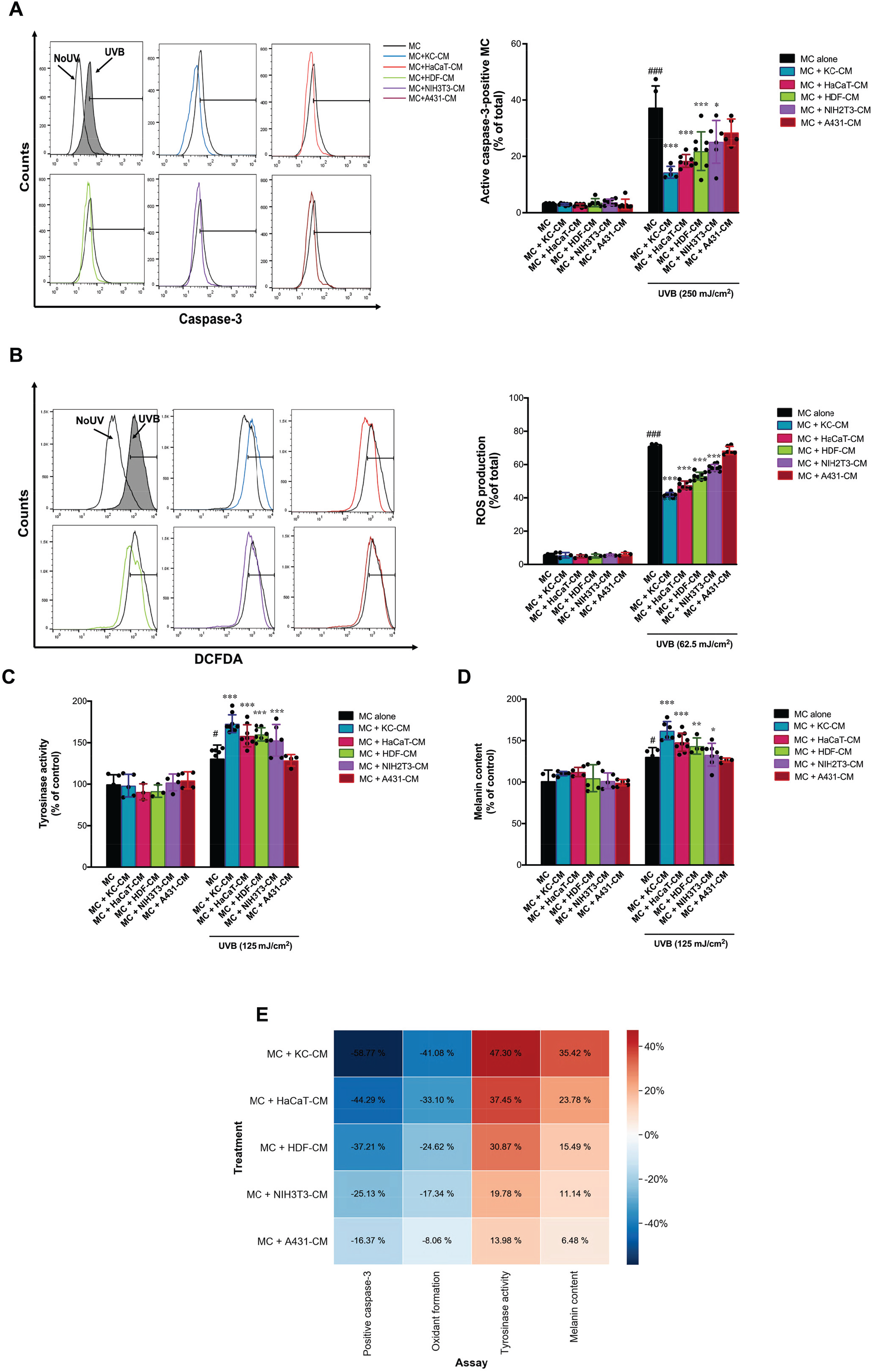
The protective effects of KC, HaCaT, HDF, NIHT3T, A431 cells on UVB-induced apoptosis, ROS formation and melanogenesis in MC treated with conditioned (CM) from KC, HaCaT, HDF, NIHT3T, A431 cells. The effects of UVB on caspase3 activation (A), ROS formation (B), melanin content (C) and tysosinase activity (D) in MC pretreated with CM from KC, HaCaT, HDF, NIHT3T, A431 irradiated with UVB (125 mJ/cm^2^) for 30 min. The statistical significance of differences between UVB-irradiated MC and UVB-irradiated MC+KC-CM, MC+HaCaT-CM, MC+HDF-CM, MC+NIH3T3-CM and MC+A431-CM was evaluated by one-way ANOVA followed by Dunnett’s test (\**P* < 0.05; \*\**P* < 0.01; \*\*\**P* < 0.001 versus UVB-irradiated MC). Heat map representing color-coded expression levels of the paracrine protective effects of KC, HaCaT, HDF, NIHT3T, A431 cells on UVB-induced MC responses (E). The red color represented the inhibitory action of paracrine protective effects. The gray color represented the activating action of paracrine protective effects.

### Candidate genes encoding paracrine factors in KC responsible for their paracine modulatory effects on UVB-mediated MC responses

We next attempted to examine the identity of paracrine factors that underlie the UVB-mediated MC response. After UVB irradiation, whole transcriptomic profiling from all four dermal cell types was profiled for unbiased exploration of all candidate paracrine factors. To ensure the correct timing for profiling gene expression changes, we first assessed gene expression changes of known paracrine factors from UVB-irradiated KC. Specifically, the expression of CRH, CRHR1, ET-1 and POMC genes were measured in KC using RT-PCR at 2, 4, 6, 8 and 12 hours post UVB irradiation (125 mJ/cm^2^) (Appendix Fig S2A). In general, we found that these secreted factors from KC reached the maximal expression changes at 6 hours post-irradiation. We then compared the expression of these paracrine factors in different dermal cell sources (KC, HaCaT, HDF and A431 cells). KC showed the most significant increases in CRHR1, ET-1 and POMC, while HaCaT only showed an increase in ET-1. On the contrary, HDF and A431 showed a negligible change in these paracrine factors (Appendix Fig S2B).

Having identified that 6 hours post UVB irradiation is the best time point for measuring the gene expression changes, we then performed whole transcriptomic profiling of KC and HaCaT cells. We performed differential gene expression (DEGs) between the UVB-irradiated and unirradiated KC and HaCaT cells. Both up-regulated and downregulated genes were identified, giving a total of 1730 genes that were identified as being differentially expressed (FDR≤0.05) (Table SI). We then shortlisted only the genes which encode secreted proteins, both inflammatory mediators and cytokines (Straussman *et al*, 2012), leaving only 46 genes of secreted factors that showed significant differential gene expressional change from the unirradiated KC and HaCAT cells (Fig 2A and B). Out of these, 16 genes were found to be upregulated in both KC and HaCaT cells (Fig 2C). The top-7 paracrine factor genes that are found to be most strongly upregulated in UVB-irradiated KC include CCL20, TNFRSF10D, CSF3 (G-CSF), IL6, IL36G, CXCL2, CXCL8 (Fig 2D). These candidate genes were then our main focus for further validation on their paracrine modulatory effects on UVB-mediated MC responses.

**Figure 2.**
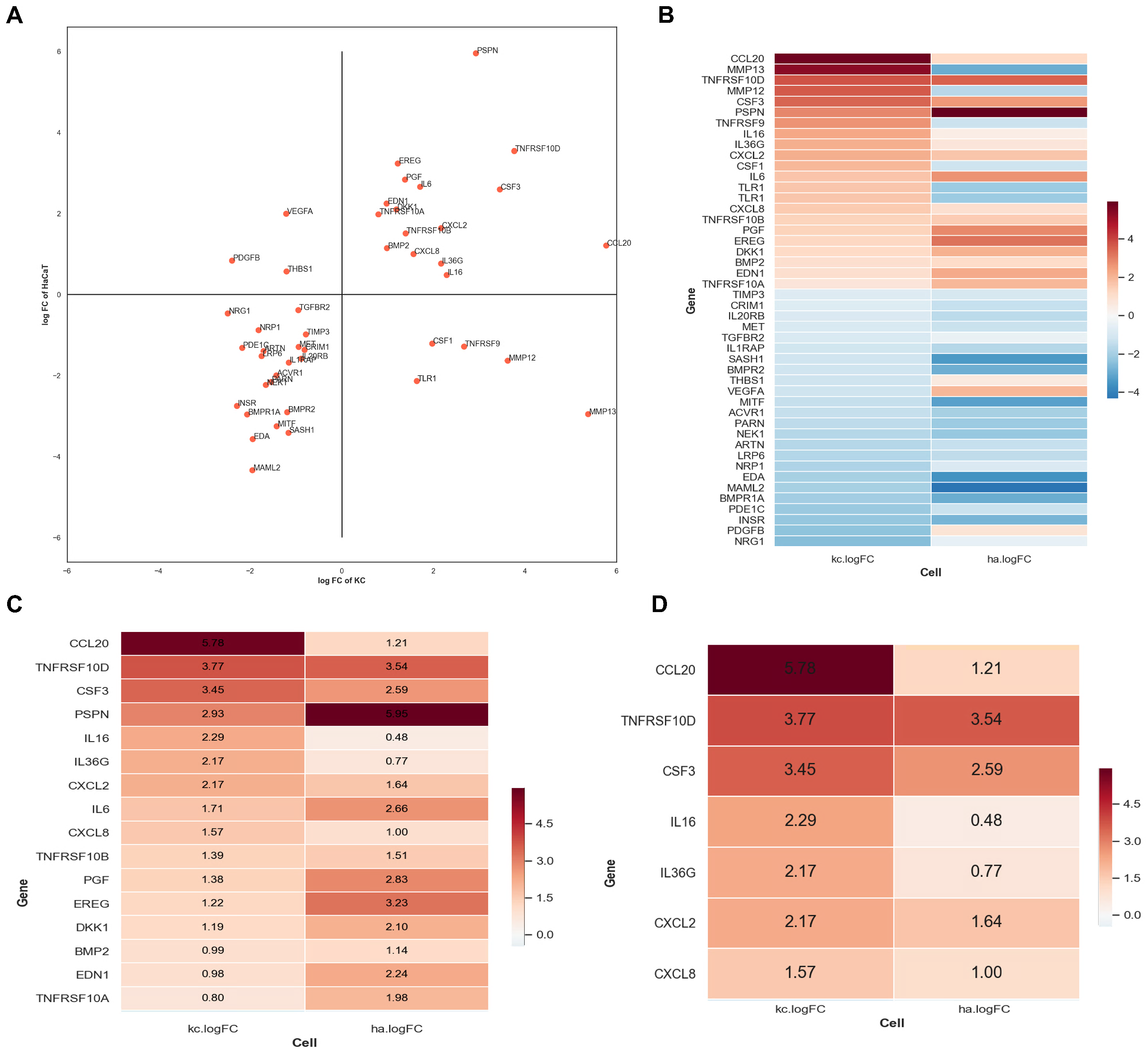
The effects of UVB on genes encoding secreted paracrine factors expression in KC and HaCaT cells. The relative differentially expressed genes of UVB irradiated keratinocytes (KC) and HaCaT (ha) cells were calculated (A). The data were presented on a log2 fold change of UVB-irradiated cells versus unirradiated cells (control cells). The x axis shows relative differentially expressed genes of UVB irradiated keratinocytes (KC), whereas the y axis shows differentially expressed genes of UVB irradiated HaCaT cells. Heatmap analysis of differentially expressed genes of UVB irradiated keratinocytes (KC) and HaCaT (ha) cells (B). The data were presented on a log2 fold change of UVB-irradiated cells versus unirradiated cells (control cells). Red color represents up-regulated genes, blue color represents non-differentially expressed genes. A total of 1730 genes were identified as being differentially expressed (FDR≤0.05) between UVB treated and control cells. UVB treated KC and HaCaT cells caused a significant expression of 46 paracrine factor genes compared to unirradiated cells (B). UVB (125 mJ/cm^2^) treatment caused 16 up-regulated transcripts in both KC and HaCaT (C). 7 paracrine factor genes were significantly upregulated in UVB-treated KC more than UVB-treated HaCaT cells (D).

### Role of the candidate paracrine factors in modulating UVB-induced apoptosis, ROS formation and melanogenesis in MC

To confirm that our identified candidate paracrine factors indeed play roles in minimizing stress responses of MC to UVB irradiation, we assessed the ability of each candidate paracrine factor to modulate UVB-mediated stress responses of MC using recombinant paracrine factors. Specifically, we incubated the MC with each of the 14 candidate paracrine factors (CSF3, CCL20, ET-1, PIGF, CXCL8, TNFRSF10B, CXCL2, IL-6, IL-16, BMP2, PSPN, IL36G, EREG, DKK1) prior to UVB irradiation. Interestingly, we found that 6 of these ligands, including CSF3 (0.3-9 nM), CCL20 (0.3-9 nM), ET-1 (0.3-9 nM), PIGF (0.3-9 nM), CXCL8 (0.3-9 nM), TNFRSF10B (1-9 nM) were able to suppress caspase-3 activation in MC (Appendix Fig S3A). Similarly, ROS formation in UVB irradiated MC was reduced with the addition of CSF3, CCL20, ET-1, CXCL8, TNFRSF10B, or BMP2 (0.3-9 nM for all six ligands) (Appendix Fig S3B). Finally, melanin content in UVB-irradiated MC was also found to be stimulated with the addition of CSF3 (1-9 nM), CCL20 (3-9 nM), ET1 (1-9 nM), PIGF (0.3-9 nM), or CXCL8 (9 nM) (Appendix Fig S3C). The heatmap analysis was also performed to summarize the modulation of UVB-induced MC responses by different recombinant proteins (Fig 3A-C). Most importantly, we found that three ligands, including CSF3, CCL20, and ET-1, could consistently suppress caspase-3 activation and ROS formation and stimulate melanin content in UVB-irradiated MC.

**Figure 3.**
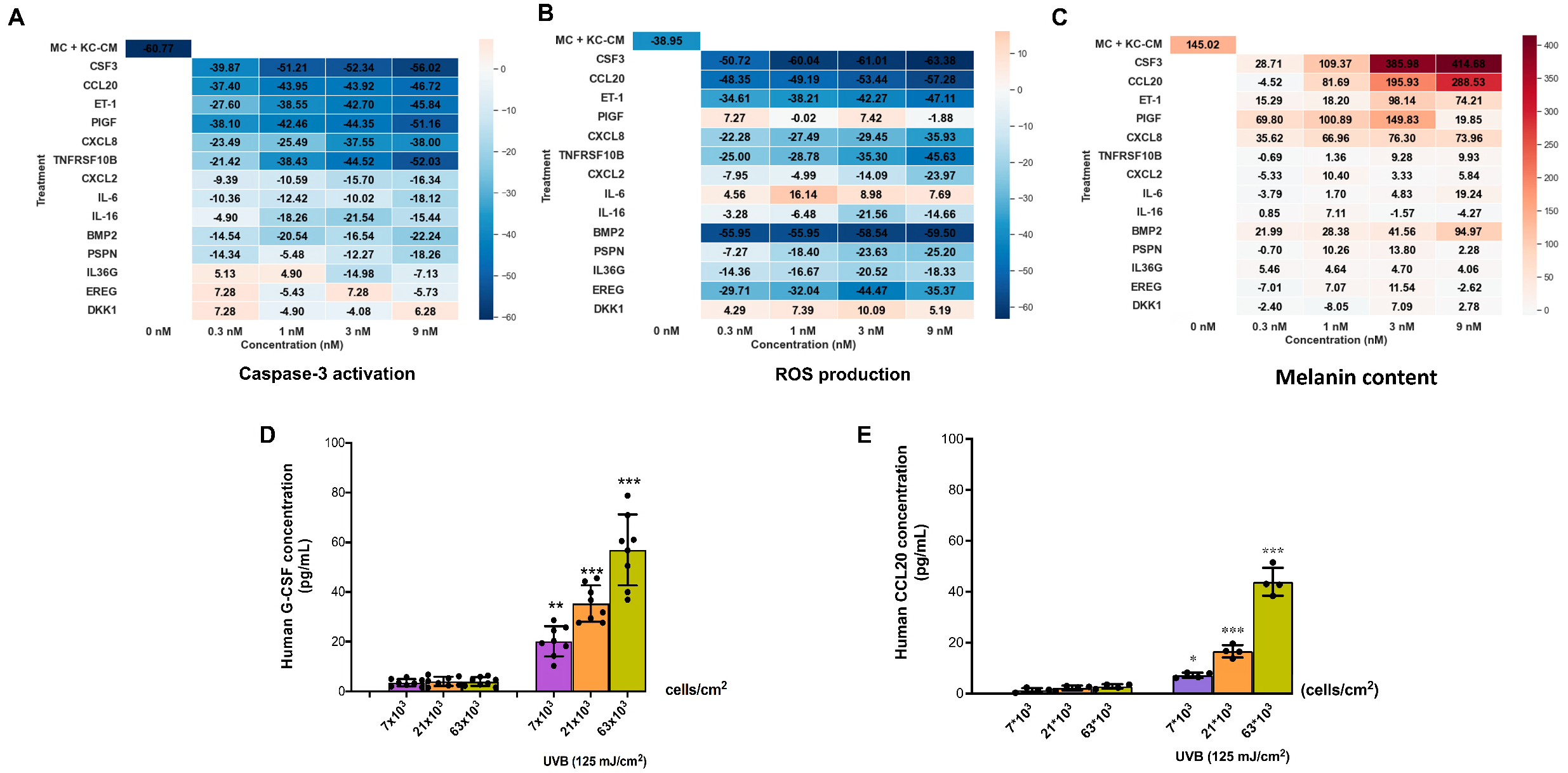
The protective effects of paracrine factors on UVB-induced apoptosis, ROS formation and melanogenesis in MC cells. Heat map representing color-coded expression levels of the paracrine protective effects of KC and 14 recombinant paracrine factors on UVB-induced apoptosis (A), ROS production (B) and melanin content (C) in MC cells. The blue color represented the inhibitory action of paracrine protective effects. The red color represented the activating action of paracrine protective effects. The effects of UVB (125 mJ/cm^2^) on G-CSF and CCL20 levels in KC cells. At 12 h post-irradiation, CM from KC at three cell concentrations (7×10^3^, 21×10^3^, 63×10^3^ cells/cm^2^) were collected and then G-CSF (D) and CCL20 (E) concentrations were measured. Data was expressed as mean ± SD. The statistical significance of differences between UVB-irradiated KC at different cell concentrations (7×10^3^, 21×10^3^, 63×10^3^ cells/cm^2^at cell ratios was evaluated by one-way ANOVA followed by Dunnett’s test (\*\**P* < 0.01; \*\*\**P* < 0.001 versus unirradiated control KC). ## *P* < 0.01; ### *P* < 0.001versus UVB-irradiated KC at cell concentrations (7×10^3^ and 21×10^3^).

We finally confirmed whether UVB irradiation induced the secretion of G-CSF and CCL20 in KC. At 12 h post-irradiation, exposure of KC at different cell plating densities (7×10^3^, 21×10^3^, 63×10^3^ cells/cm^2^) to UVB (125 mJ/cm^2^) led to a corresponding increase of both G-CSF (20.1 ± 5.7, 36.92 ± 6.3, 58.26 ± 10.5 pM) and CCL20 (7.12 ± 1, 16.61 ± 2.4, 43.90 ± 5.5 pM) (Fig 3D and E).

### The modulatory effects of the candidate paracrine factors on UVB-induced upregulation of melanogenesis-related genes (tyrosinase and TRP-1) in MC cells

Our results also revealed that G-CSF and CCL20 could upregulate major melanogenesis-related genes including tyrosinase and TRP-1, in UVB-irradiated MC, indicating the involvement of G-CSF and CCL20 in the paracrine actions of KC on the melanogenic response of MC to UVB. As observed in our study, among several UVB-responsive genes involved in paracrine signaling identified in KC, the paracrine factors, G-CSF and CCL20, might play a crucial role in KC’s paracrine action against UVB-induced stress responses of MC.

The previous results indicated that paracrine factor ligands, including G-CSF and CCL20, provide the strongest effects on MC in response to UVB-induced melanogenesis (Melanin content). Then we next investigate the effects of recombinant human G-CSF and CCL20 on crucial melanogenesis-related genes, including tyrosinase and TRP-1 (Nguyen N.T and Fisher D.E, 2019). Our results revealed that pretreatment of MC with the recombinant G-CSF and CCL20 (3 nM) resulted in a significant induction of tyrosinase (Fig 4A) and TRP1 mRNA expression (Fig 4B) in UVB-irradiated MC. The upregulation of tyrosinase and TRP-1 mRNA was also observed in MC treated with KC-CM and the positive control, α-MSH (200 nM), an effective activator of melanogenesis-related genes including tyrosinase and TRP-1 genes.

**Figure 4.**
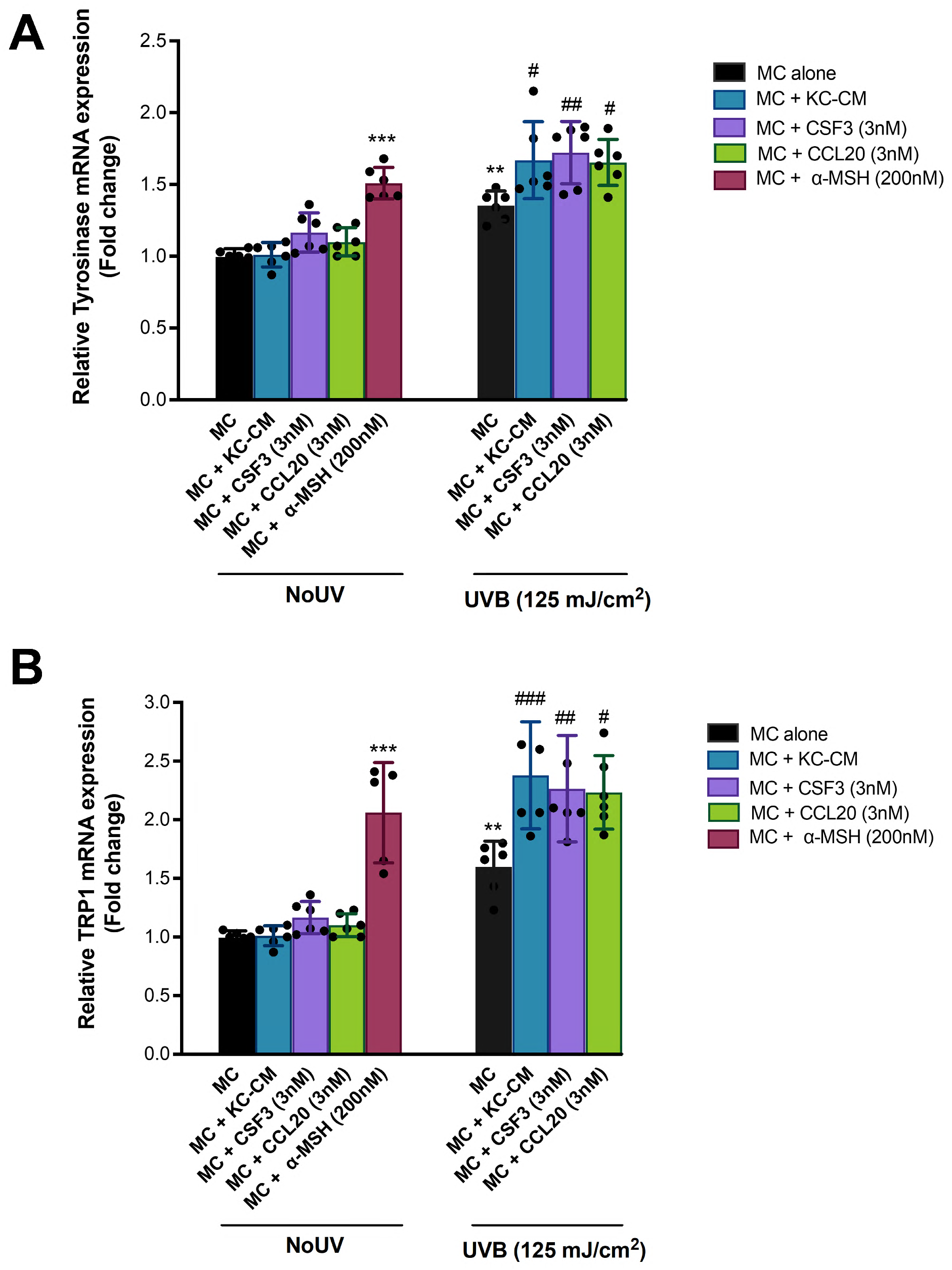
The protective effects of the paracrine factors on UVB-induced transcriptional activation of MITF in MC cells. The effects of UVB (125 mJ/cm^2^) on tyrosinase (A) and TRP1 (B) mRNA expression in MC pretreated with CM from KC, irradiated with UVB (125 mJ/cm^2^), G-CSF, CCL20 (9 nM) and œ-MSH (200 nM). MC were harvested at 1 h after UVB irradiation for determination of ROS formation. Data was expressed as mean ± SD. T he statistical significance of differences was evaluated by one-way ANOVA followed by Dunnett’s test. \*\**P* < 0.01; \*\*\**P* < 0.001 versus unirradiated cells. The statistical significance of differences between UVB-irradiated MC and UVB-irradiated MC+KC-CM, and G-CSF and CCL20 was evaluated by one-way ANOVA followed by Dunnett’s test (#*P* < 0.05; ##*P* < 0.01; ###*P* < 0.001 versus UVB-irradiated MC).

### The modulatory effects of the recombinant paracrine factors on UVB-induced melanogenesis in mouse skin exposed to UVB irradiation

Since transcriptional profiling of KC in the skin can be influenced by the surrounding non-KC cell types (Gazel *et al*, 2003), we next asked whether the observed effect of KC-originated paracrine factors can be observed in the physiological context. We used mouse models with UVB-induced photodamage of the skin to explore the intercellular interactions between KC and MC. H&E stained sections of mouse skin tissues revealed that UVB irradiation (500, 750, 1,000 mJ/cm^2^) caused a substantial and dose-dependent increase in epidermal thickness (11.90 ± 1, 14.99 ± 0.6 and 12.83 ± 0.7 uM, respectively) (Fig 5A and B). With the exposure of UVB (750 mJ/cm^2^), we observed expression of the G-CSF and CCL20 in KC (Appendix Fig S4). By profiling the abundance of G-CSF and CCL20 proteins based on immunofluorescent staining with increasing UVB dosing, we revealed a correlative increase of both proteins with the melanogenic responses to UVB irradiation in mouse skin (Fig 5C-G). Specifically, UVB irradiation (500, 750, 1,000 mJ/cm^2^) led to a significant dose-dependent induction of protein expressions of G-CSF (133.27 ± 11.7, 140.68 ± 21.2 and 125.11 ± 17.6% of control, respectively) and CCL20 (144.36 ± 30.6, 162.07 ± 22.4 and 135.87 ± 20.4% of control, respectively) (Fig 5C-F) as well as tyrosinase (185.72 ± 23, 144.70 ± 22.8 and 128.29 ± 16.9 % of control, respectively) (Fig 5C, E and G). Therefore, our observations suggested that expressions of G-CSF and CCL20 protein in the epidermis positively correlated with the expression of tyrosinase, the enzyme responsible for melanin synthesis.

**Figure 5.**
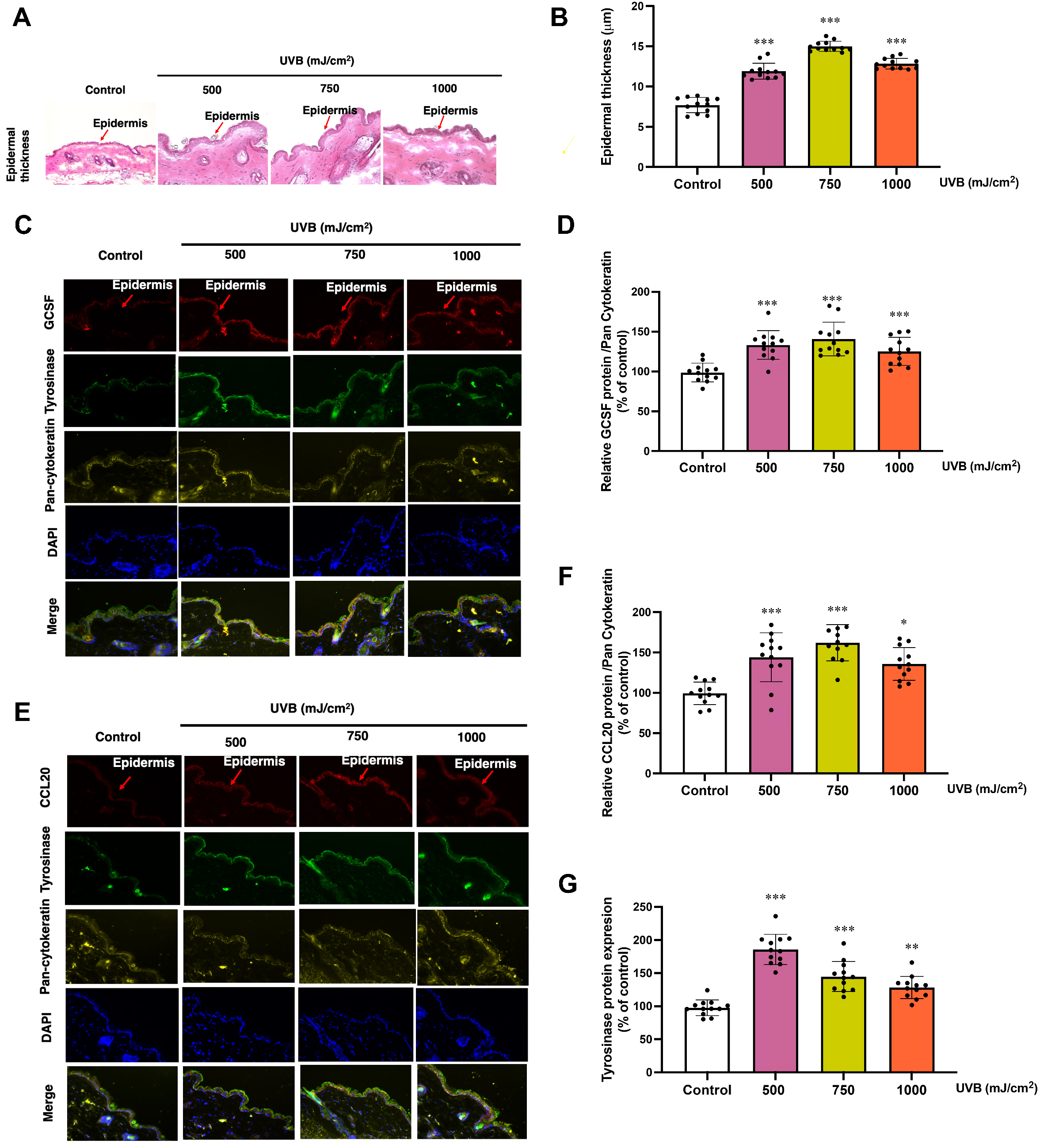
Epidermal thickness and the expression of G-CSF, CCL20 and tyrosinase protein in mouse skin exposed to UVB irradiation. Epidermal thickness (A), the immunofluorescence of G-CSF (B), CCL20 (C), Tyrosinase (B and C) and pan-cytokeratin which is keratinocyte marker (B and C) were collected at 12 h following the final UVB exposure. The summary graph with the statistical analysis of the epidermal thickness (D) and the relative protein levels of GCSF (E), CCL20 (F) to pan-cytokeratin and tyrosinase (G) were quantified by ImageJ and graphpad prism software and were expressed as mean ± SD, 1 N (dot) = one area from 4 areas in a mouse. A mouse provides 3 N. **\*** *P* < 0.05, \*\**P* < 0.01, \*\*\**P* < 0.001 versus non-irradiated group by Student’s *t*-test.

## Discussion

KC and MC are the major target cells of UVB radiation in the skin and the microenvironment created by KC potentially influences the survival and function of MC in response to various exogenous insults including UVB (Sample & He, 2018). Our previous findings and other studies suggested that KC can protect against UVB-mediated photodamage in MC by reducing apoptosis, oxidative stress, DNA damage, and inflammatory responses and promoting melanogenesis (Jeayeng *et al*, 2017; Upadhyay PR *et al*, 2017). To gain further insight into the crosstalks between MC and the microenvironment created by KC, which is critical in governing epidermal homeostasis, we aimed in this study to identify the underlying paracrine factors from KC that play key roles in protecting MC from the UVB-induced photodamage. Having identified different dermal cell sources with differential protective effects against UVB-mediated photodamage in MC, we reasoned that the expression level of the underlying paracrine factors in KC must show corresponding expressional levels in these different dermal cell sources. This strategy identified major paracrine factor genes, including CCL20, TNFRSF10D, CSF3, IL6, IL36G, CXCL2, and CXCL8. By, we showed that adding these recombinant paracrine factors (at least, G-CSF, CCL20, ET1, PIGF, CXCL8) can dose-dependently suppress caspase-3 activation and ROS formation as well as stimulated melanin content in UVB-irradiated MC. G-CSF and CCL20 were confirmed to be secreted from the KC-derived CM after UVB exposure. With the mouse skin model, we also proved that CSF3 and CCL20 protein abundance near KC increased with the elevated UVB dosage.

While epidermal KC and dermal fibroblasts can regulate the growth, differentiation, and pigmentation of MC in a paracrine fashion (Wang *et al*, 2017), our findings suggested that the microenvironment created by KC might primarily influence phenotypic and functional responses of MC to UVB. Since MC act as stress sensors in the epidermis vulnerable to numerous environmental threats (Slominski *et al*, 2004), biological and physiological regulation of MC by KC could therefore serve as a photoprotective signature and a potential therapeutic target for UVB-mediated photodamage. In fact, several soluble paracrine factors, e.g., hepatocyte growth factor (HGF), Neuregulin-1 (NRG-1), transforming growth factor-b (TGFb), and basic fibroblast growth factor (bFGF) can be produced by KC and HDF and were shown to regulate MC homeostasis (Wang *et al*, 2017). The commonly known UVB signature genes, such as ET-1 and POMC, were suggested to be the crucial KC-derived paracrine factors regulating MC homeostasis and function in response to UVR (Zimmermann *et al*, 2001; Hyter *et al*, 2013; Rousseau *et al*, 2007). In our study, expression levels of the seven upregulated transcripts identified as the UVB-responsive gene set (6 h post-UVB) were higher than those of the commonly known UVB signature genes previously reported. Furthermore, our observations indicated that, at an early time point (6 h after exposure), UVB at a high dose (125 mJ/cm^2^) that drive stress responses, including apoptosis, ROS formation and melanogenesis in MC, could regulate various genes encoding secreted paracrine factors in KC but did not significantly affect genes involved in oxidative stress/antioxidant responses, which were primarily modulated by UVA irradiation (Sesto *et al*, 2002). Our findings are also in agreement with other previous studies suggesting that, in response to UVB exposure, most transcripts (>80%) in KC were downregulated at early time points (4-6 h post-UVB) (Sun X *et al*, 2016) and the upregulated transcripts included genes encoding the secreted paracrine factors including growth factors and cytokines (López-Camarillo C *et al*, 2012).

Previous studies suggested that colony-stimulating factors, including CSF2 (GM-CSF) and CSF3 (G-CSF), and CCL20, are paracrine factors involved in the regulation of the cutaneous homeostasis, tissue repair and regeneration, hyperpigmentation, inflammation as well as immune responses (Feliciani *et al*, 1996; Kennedy *et al*, 2012). G-CSF was observed to improve wound healing in mice via anti-apoptotic and anti-inflammatory actions (Huang *et al*, 2017). A previous study using models of UVB-induced epidermal injury suggested that an upregulation of cytokine CCL20, observed as an early response to injury, was involved in the early stages of wound healing (Kennedy *et al*, 2012). Moreover, GM-CSF derived from KC was demonstrated to play a role in controlling melanogenesis in UVA-irradiated MC (Imokawa *et al*, 1996), and the proliferation and differentiation of mouse MC from UVB-induced pigmented spots (Hirobe *et al*, 2002). Our results revealed that G-CSF and CCL20 could upregulate major melanogenesis-related genes including tyrosinase and TRP-1, in UVB-irradiated MC, indicating the involvement of G-CSF and CCL20 in the paracrine actions of KC on the melanogenic response of MC to UVB. As observed in our study, among several UVB-responsive genes involved in paracrine signaling identified in KC, the paracrine factors, G-CSF and CCL20, might play a crucial role in KC’s paracrine action against UVB-induced stress responses of MC. Further study is still required to elucidate the complex signaling pathways involved in the regulation of crosstalk between KC and MC to give insights into potential biomarkers for the development of targeted prevention and treatment of UVB-induced skin conditions related to MC damage.

Our study also inferred that the upregulated transcripts might be involved in the skin defense response to UVB-mediated acute injury of the skin. High-throughput approaches have been employed to identify UV-responsive genes and proteins in cultured skin cells and mouse skin (Sun *et al*, 2016; Wang *et al*, 2019). Additionally, the transcriptional responses to UVR are complex, depending on wavelengths and intensity of UV as well as time after the irradiation. Some regulated transcripts have several biological functions and associate with multiple pathways. Since the roles of UV-responsive genes in KC involve regulations of different cellular functions and cell fates including apoptosis, cell proliferation, and growth (Shen *et al*, 2016), it should be taken into account that paracrine factors (including growth factors and cytokines) act as double-edged swords because they could either prevent premature cell death and promote tissue repair or contribute to immune responses, inflammation, and carcinogenesis (Mueller *et al*, 1999). In response to KC environmental insults, the balance of anti-apoptotic and pro-apoptotic cascades must be tightly controlled to maintain the balance between cell survival and death (Raj *et al*, 2006). Hence, further study is required to evaluate clinical relevance and validate the specificity of the UV signature genes associated with an increased risk for skin photodamage and photocarcinogenesis.

In summary, our study demonstrated that the G-CSF and CCL20 are among the major paracrine factors secreted by KC that contribute to minimizing MC’s stress responses to UVB. The UVB-responsive genes in KC observed in our study may be promising in developing clinical UVB biomarkers for predicting the skin’s susceptibility to photodamage. Furthermore, the UVB signature paracrine factors might be exploited as targets to develop novel and effective mechanism-based approaches to prevent and treat skin photodamage.

## Materials and Methods

### Cell culture

Primary human epidermal melanocytes (MC) and primary human epidermal keratinocytes (KC) were obtained from Invitrogen (NY, USA). MC was cultured in Medium

254 (#M-254-500) supplemented with human melanocyte growth supplement (HMGS) according to the manufacturer’s instructions. Primary human epidermal keratinocytes (KC) were cultured in Medium 154 (#M-154CF-500) supplemented with human keratinocyte growth supplement (HKGS). Human keratinocyte (HaCaT) cells (Cell Lines Service, Heidelberg, Germany) were grown in high glucose (4.5 g/L) Dulbecco’s modified Eagle’s medium (DMEM) and Ham’s F-12 (DMEM/F-12) supplemented with 10% fetal bovine serum and 1% penicillin (100 U/ml)/streptomycin (100 mg/ml). Primary human dermal fibroblast (HDF) was grown in high glucose (4.5 g/L) Dulbecco’s modified Eagle’s medium (DMEM) supplemented with 10% fetal bovine serum, 1% 200 mM glutamax and 1% Antibiotic-Antimycotic Solution (Mediatech, Inc. Manassas, VA). Primary mouse embryonic fibroblast cells (NIH3T3) and epidermoid carcinoma (A431) were cultured in Dulbecco’s minimal essential media (DMEM) supplemented with 10% fetal bovine serum and 1% Antibiotic-Antimycotic Solution (Mediatech, Inc. Manassas, VA). All cells were maintained at 37 °C in a humidified air of 5% CO2 (PCO2 = 40 Torr) (a Forma Scientific CO2 Water Jacketed Incubator).

### Preparation of cell-derived conditioned medium (CM)

KC, HaCaT, HDF, NIH3T3 and A431 were seeded at a density of 5 × 10^5^ cells/well in 6 cm^2^ dishes. Then CM was prepared from a cell density of 21×10^3^ for all experiments to determine the effects of CM on apoptosis, ROS formation and melanogenesis in MC. HDF, NIH3T3 and A431 supernatants were used as non-keratinocyte groups to compare the effects of paracrine factors between keratinocyte and non-keratinocyte cells. Conditioned-KC, HaCat, HDF, NIH3T3, and A431 supernatants were prepared by irradiation of 5 skin cells in DPBS with UVB (125 mJ/cm^2^), and the DPBS will be changed to DMEM medium and collected at 12 h post-irradiation and used as KC-CM, HaCat-CM, HDF-CM, NIH3T3-CM and A431-CM for treatment of MC. MC was pre-incubated with KC-CM, HaCat-CM, HDF-CM, NIH3T3-CM and A431-CM for 30 min, subjected to UVB irradiation and harvested at one h following 62.5 mJ/cm^2^ UVB irradiation for determination of ROS formation, 12 h after 250 mJ/cm^2^ irradiation for caspase-3 activation and 12 h after 125 mJ/cm^2^ irradiation for melanin content, tyrosinase activity.

### UVB irradiation

MC, KC, HaCaT, HDF, NIH3T3 and A431 cells were irradiated with UVB under a thin layer of Dulbecco’s phosphate-buffered saline (DPBS). The UV intensity determined at a distance of 21 cm from the UVR lamp was 1 W/cm2 using a UV meter (Dr. Honle, Martinsried, Germany). Following our previous observations, the culture plates were exposed for 22.5, 45 s or 1 min 30 s to achieve a single dose of 62.5, 125, or 250 mJ/cm2, respectively (Jeayeng *et al*, 2017). Immediately after UVB exposure, DPBS were replaced with Medium 254 without HMGS for MC and DMEM without FBS for KC, HaCaT, HDF, NIH3T3, and A431. MC was harvested at one h post-irradiation for ROS formation and 12 h post-irradiation for apoptosis using cleaved caspase-3 assays, melanin content, and tyrosinase activity. KC, HaCaT, and HDF cells were harvested at six h post-irradiation for mRNA expression.

### Determination of intracellular oxidant formation by flow cytometry

DCFH-DA was used to determine intracellular reactive oxygen species (ROS). The principle of the DCFH-DA method is detecting the intensity of green fluorescence that occur after deacetylation and subsequent oxidation product of the probe. DCFH-DA gets into cells and accumulates mainly in the cytosol. Then it is hydrolyzed in the cells to DCFH, further oxidized by oxidants (e.g., H_2_O_2_) to fluorescent 2,7-dichlorofluorescein (DCF). Thus, the fluorescence reflected the overall oxidative stress and oxidant formation in cells which can be visualized and detected using FACS Calibur.

The assay is based on the conversion of non-fluorescent dichlorofluorescein (H2DCFDA) to the fluorescent 2,7-DCF upon oxidation by intracellular ROS. After one hour of UVB irradiation, MC was washed. Then, cells were incubated in DPBS with 20 *μ*M H_2_DCFDA at 37 °C for 30 min and analyzed by flow cytometry using a fluorescence-activated cell sorter (FACS-Calibur) as previously described (Lohakul *et al*, 2021).

### Measurement of active caspase-3

Caspase-3 is a crucial mediator of programmed cell death (apoptosis). In the apoptotic cell, caspase3 can be activated by extrinsic (death ligand) and intrinsic (mitochondrial) pathways. In cells undergoing apoptosis, caspase-3 is an essential protease activated during the early stages of apoptosis and synthesized as an inactive pro-enzyme.

Active caspase-3 was measured using PE Active Caspase-3 Apoptosis Kit (BD Biosciences, USA) according to the manufacturer’s instructions. After 12 hours of UVB irradiation, MC was washed to determine active caspase-3. Briefly, MC was fixed and permeabilized using the Cytofix/CytopermTM for 30 min, pelleted, and washed with Perm/Wash TM Buffer. Cells were subsequently stained with the rabbit anti-active caspase-3 antibody (clone C92-605) (BD Biosciences, USA) in the dark. Cells were then washed and resuspended in Perm/WashTM Buffer and analyzed by flow cytometry as described in our previous report (Jeayeng *et al*, 2017).

### Determination of melanin content and tyrosinase activity

Melanin content and tyrosinase activity were determined in MC. Cells harvested at 12 hr -postirradiation. The melanin and tyrosinase activity monitored by dopachrome formation was measured spectrophotometrically at 475 by a spectrophotometer. The amount of melanin (µg/mg protein) was calculated by comparing it with a standard curve generated using synthetic melanin. The tyrosinase activity (unit/mg protein) was calculated by comparison to a standard curve using tyrosinase (2034 U/mg) as previously described (Chaiprasongsuk *et al*, 2016).

### Quantitative real-time reverse transcriptase-polymerase chain reaction (RT-PCR) for determination of mRNA expression

mRNA levels including CRH, CRHR1, ET-1 and POMC were determined by RT-PCR. Total RNA was isolated using the Illustra RNAspin Mini RNA Isolation Kit (GE Healthcare, UK), and reverse transcription was carried out using the Improm-II reverse transcriptase (Promega, Medison, USA) under the conditions described in the kit manual. Primers for PCR were designed using the Primer Express software version 3.0 (Applied Biosystems, USA). Sequences of PCR primer (in 5’→3’ direction) were as follows: CRH (product sizes = 144 bp) sense, CTCCGGGAAGTCTTGGAAAT, and antisense, GTTGCTGTGAGCTTGCTGTG; CRHR1 (product sizes = 100 bp) sense, TGGATGTTCATCTGCATTGG, and antisense, TGCCAAACCAGCACTTCTC; ET-1 (product sizes = 274 bp) sense, TCTACTTCTGCCACCTGGAC, and antisense, CACTTCTTTCCCAACTTGGAAC; POMC, exon 3 (product sizes = 152 bp) sense, AGCCTCAGCCTGCCTGGAA, and antisense, CAGCAGGTTGCTTTCCGTGGTG. The mRNA level was calculated by normalizing with the expression level of GAPDH mRNA. The mean Ct from mRNA expression in cDNA from each sample was compared with the mean Ct from GAPDH determinations from the same cDNA samples as described in our previous report (Jeayeng *et al*, 2017).

### RNA sequencing analysis (RNA-seq)

RNA was extracted using Trizol, purified by Ambion RNA isolation kit (Invitrogen) and treat with PureLink DNase set (Invitrogen). RNA samples were quantified by measuring absorbance at 260nm with the Qubit RNA HS assay kit. The integrity of RNA was analyzed by running on agarose gel electrophoresis and measuring RIN value with Agilent 2100 Bioanalyzer (Agilent Technologies). All samples showed high RNA integrity with intact 28sRNA, intact 16sRNa, and RNA integrity number (RIN)>8.

The total RNA of each sample was quantified and qualified by Agilent 2100 Bioanalyzer (Agilent Technologies, Palo Alto, CA, USA), NanoDrop (Thermo Fisher Scientific Inc.), and 1% agarose gel. One μg total RNA with the RIN value above seven was used for the following library preparation.

Next-generation sequencing library preparations were constructed according to the manufacturer’s protocol (NEBNext® Ultra™ RNA Library Prep Kit for Illumina®). The poly(A) mRNA isolation was performed using NEBNext Poly(A) mRNA Magnetic Isolation Module (NEB). The mRNA fragmentation and priming were performed using NEBNext First Strand Synthesis Reaction Buffer and NEBNext Random Primers. First-strand cDNA was synthesized using ProtoScript II Reverse Transcriptase, and the second-strand cDNA was synthesized using Second Strand Synthesis Enzyme Mix. The purified (by AxyPrep Mag PCR Clean-up (Axygen)) double-stranded cDNA was then treated with End Prep Enzyme Mix to repair both ends and add a dA-tailing in one reaction, followed by a T-A ligation to add adaptors to both ends. Size selection of Adaptor-ligated DNA was then performed using AxyPrep Mag PCR Clean-up (Axygen), and fragments of ∼360 bp (with the approximate insert size of 300 bp) were recovered. Each sample was then amplified by PCR for 11 cycles using P5 and P7 primers, with both primers carrying sequences that can anneal with flow cell to perform bridge PCR and P7 primer having a six-base index allowing for multiplexing. The PCR products were cleaned up using AxyPrep Mag PCR Clean-up (Axygen), validated using an Agilent 2100 Bioanalyzer (Agilent Technologies, Palo Alto, CA, USA), and quantified by Qubit 2.0 Fluorometer (Invitrogen, Carlsbad, CA, USA).

Then libraries with different indices were multiplexed and loaded on an Illumina HiSeq instrument according to the manufacturer’s instructions (Illumina, San Diego, CA, USA). Sequencing was carried out using a 2×150bp paired-end (PE) configuration; image analysis and base calling were conducted by the HiSeq Control Software (HCS) + OLB + GAPipeline-1.6 (Illumina) on the HiSeq instrument

Sequencing was performed on the Illumina HiSeq platform, in a 2×150bp paired-end (PE) configuration, with 2.0 Gb raw data per sample. Base-calling is performed by Illumina RTA software in a sequencer, and further demultiplexing is performed by Illumina bcl2fastq 2.17 software based on the index information. The number of reads and quality score (Q30) are also counted.

Alignment of reads to the human reference (hg19) was done using STAR version v2.5.3a (Dobin A *et al*, 2013 on the Basepair website (https://www.basepairtech.com/). Other pipelines were used, including ‘sambamba’ (v0.6.6) (Tarasov A *et al*, 2015) for sorting the BAM files, ‘samtools’ (v1.6) (Li H *et al*, 2009) for indexing the BAM files, ‘fastp’ (v0.19.4) (*Chen S et al*, 2018) for the read trimming and QC and ‘subread’ (v1.6.2) (Liao Y *et al*, 2013) for the gene/isoform quantification. Trimming of low-quality base pairs was done using Phred quality score (Q score) at ten as the cut-off. (Williams CR *et al*, 2016) Differential gene expression was done using the BioJupies platform (https://biojupies.cloud) (Torre D *et al*, 2018). Significant level was considered if the adjusted p-value < 0.05.

### Determination of G-CSF and CCL20 levels by ELISA

G-CSF and CCL20 levels in culture supernatants were determined using competitive enzyme immunoassay kits from DCS50 and DM3A00, R&D Systems, Inc., USA, according to the manufacturer’s instructions. Sample or G-CSF, CCL20 standards were added to the immuno plate pre-coated with secondary antibody. An anti-G-CSF CCL20 antibody was added, followed by a biotinylated peptide. The biotinylated peptide then interacted with streptavidin-horseradish peroxidase (HRP), which catalyzed the substrate solution.

### Determination of UVB-induced cell damage *in vivo*

All animal experiments were reviewed and approved by the Siriraj Animal Care and Use Committee (SiACUC), SiACUP 018/2562. Three-week-old female BALB/c wild-type mice were purchased from National Laboratory Animal Center, Mahidol University and maintained under controlled conditions (25 ± 2 °C with 55 ± 5% relative humidity in a 12-h light:12-h dark cycle) using an isolator caging system. Water and food diet were available ad libitum during the experimental period as previously described (Lohakul *et al*, 2021; Lohakul *et al*, 2022, Chaiprasongsuk *et al*, 2017). Mice were randomized into four groups of 4 mice. Group I (control), without UVB exposure. Group II, UVB 2 time of 250 mj/cm^2^ irradiation (a cumulative total dose of 500 mJ/cm^2^). Group III, UVB 3 time of 250 mj/cm^2^ irradiation (a cumulative total dose of 750 mJ/cm^2^). Group IV, UVB 3 time of 250 mj/cm^2^ irradiation (a cumulative total dose of 1000 mJ/cm^2^). Mice were anesthetized by intraperitoneal injection (*i*.*p*.) of 100:10 mg/kg of ketamine/xylazine cocktail. Dorsal skin was removed at 12 h after the last exposure of UVB, embedded in Tissue-Tek® OCT compound and directly snap-frozen (liquid N_2_), and stored at -80 °C until microtome sectioning. Skin thickness was assessed by hematoxylin and eosin (H&E) and immunofluorescence (IF) staining. CCL20, G-CSF, tyrosinase, pan-cytokeratin (keratinocyte marker) and gp100 (melanocyte marker) were assessed by immunofluorescence and DAPI was used to counterstain nuclei. Image analysis was performed using Image J.

### Statistical Analysis

Data are expressed as mean ± standard deviation of the mean (SD) of at least three separate experiments (n ≥ 3) performed on different days using freshly prepared reagents. Statistical significance of differences between different groups was evaluated by independent t-test (Student’s; 2 populations) or one-way analysis of variance (ANOVA) followed by Tukey or Dunnet tests, where appropriate, using Prism (GraphPad Software Inc., San Diego, CA)

## Acknowledgment

This work was supported by the Mahidol University (Fundamental Fund: fiscal year 2023 by National Science Research and Innovation Fund (NSRF), “Mahidol University” grant and

Siriraj Research and Development Fund Type II [grant number R016233017], Faculty of Medicine Siriraj Hospital, Mahidol University and Mahidol.

## Author contributions

Conceptualization, U.P. and S.S.; methodology, S.J., M.S. and P.M.; software, S.J., M.S., P.M., P.B. and P.K.; validation, S.J., M.S. and P.M.; formal analysis, S.J., M.S., P.M., P.B. and P.K.; investigation, S.J., M.S., P.M. and U.P.; resources, U.P.; data curation, S.J., M.S., P.M., P.B. and P.K.; writing—original draft preparation, S.J., S.S. and U.P.; writing— review and editing, S.J., S.S. and U.P.; visualization, U.P.; day to day supervision, U.P.; project administration, U.P.; funding acquisition, U.P. All authors have read and agreed to the published version of the manuscript.

